# Tailored polymeric, photovoltaic, and near-infrared-responsive neuroprosthesis

**DOI:** 10.1101/2020.01.27.920819

**Authors:** Marta Jole Ildelfonsa Airaghi Leccardi, Naïg Aurelia Ludmilla Chenais, Laura Ferlauto, Maciej Kawecki, Elodie Geneviève Zollinger, Diego Ghezzi

**Affiliations:** Medtronic Chair in Neuroengineering, Center for Neuroprosthetics and Institute of Bioengineering, School of Engineering, École polytechnique fédérale de Lausanne, 1202 Geneva, Switzerland; Laboratory for Nanoscale Materials Science, Empa, 8600 Dübendorf, Switzerland; Department of Physics, University of Basel, 4056 Basel, Switzerland

## Abstract

Organic materials, such as conjugated polymers, are attractive building blocks for bioelectronic interfaces. In particular, organic semiconductors showed excellent performances in light-mediated excitation and silencing of neuronal cells and tissues. However, the main challenges of these organic photovoltaic interfaces compared to inorganic prostheses are the limited stability of conjugated polymers in the aqueous environment and the exploitation of materials only responsive in the visible spectrum. In this report, we show a new photovoltaic organic interface tailored for neuronal stimulation in the near-infrared spectrum. Also, we adjusted the organic materials by chemical modification in order to improve the stability in aqueous environment and to modulate the photoelectrical stimulation efficiency. As proof of principle, we tested this interface for retinal stimulation. Our results provide an efficient, reliable, and stable implant applicable for neural stimulation.

## Introduction

Conjugated polymers and organic semiconductors have proven to be an efficient tool in bioelectronic interfaces and neuroprostheses to modulate the neuronal activity by converting light pulses into electrical or thermal stimulation^1–11^. Flexibility, lightweight, and biocompatibility are among the main advantages of using organic technology in bioelectronic interfaces. Nevertheless, compared to their inorganic counterparts, some challenges of functional organic materials remain open. First, water-induced swelling, degradation, and delamination are among the most critical aspects of organic interfaces implanted into the body. Second, in the case of photovoltaic interfaces, the electrical properties of the photovoltaic cell must be tailored to meet the desired conditions of electrical stimulation. Last, organic semiconductors used in photovoltaic bioelectronic interfaces typically have high sensitivity in the visible spectrum. However, the use of near-infrared (NIR) light is desirable because it allows a higher penetration into the tissue.

A notable example is the exploitation of conjugated polymers in photovoltaic retinal prostheses, which were proposed to induce artificial vision in blind patients^5,12–15^. Since photovoltaic retinal prostheses wirelessly convert the light entering the pupil into electrical stimuli, they act as artificial photoreceptors to activate surviving retinal cells. In the past years, conjugated polymers (*i.e.* poly(3,4-ethylenedioxythiophene)-poly(styrenesulfonate), PEDOT:PSS; regioregular poly(3-hexylthiophene-2,5-diyl), P3HT) and other organic semiconductors ([6,6]-phenyl-C61-butyric acid methyl ester, PC_60_BM) have been successfully exploited to build an organic photovoltaic subretinal interface able to improve visual acuity in dystrophic rats^5,7,13,16,17^. More recently, our group described a novel foldable and photovoltaic wide-field epiretinal prosthesis (POLYRETINA) based on the P3HT:PC_60_BM bulk heterojunction (BHJ), with the capability to activate retinal ganglion cells with short pulses of green light^12^. In the recent years, several organic photovoltaic retinal prostheses were developed based on PEDOT:PSS and P3HT:PC_60_BM BHJ; however, these materials might not possess optimal properties for photovoltaic retinal stimulation.

In humans and primates, color vision is based on three types of opsins associated to three different cones: the short (S-), the medium (M-), and the long (L-) wavelength sensitive cones having distinct but overlapping absorption spectra (**Fig. 1a**)^18^. The L-cone has the most right-shifted spectral absorbance with a maximum around 564 nm (**Fig. 1a**, circles). The absorbance spectra of the P3HT:PC_60_BM BHJ and the retinal photoreceptors largely overlap; thus, the green light used to excite previously proposed retinal interfaces based on the P3HT:PC_60_BM BHJ may not be optimal due to the possible activation of remaining cones and rods in patients with residual natural vision, such as in age-related macular degeneration. In this case, the use of NIR light is desirable to activate photovoltaic prostheses without interfering with the residual natural vision. Moreover, the irradiance levels typically required to stimulate retinal neurons with photovoltaic prostheses (hundreds of μW mm^−2^) may still be perceived (if visible light is used) even by blind patients without residual vision. Last, according to the standard for optical safety, the maximum permissible exposure (MPE) for ophthalmic applications increases in the NIR spectrum^19^.

**Figure 1.**
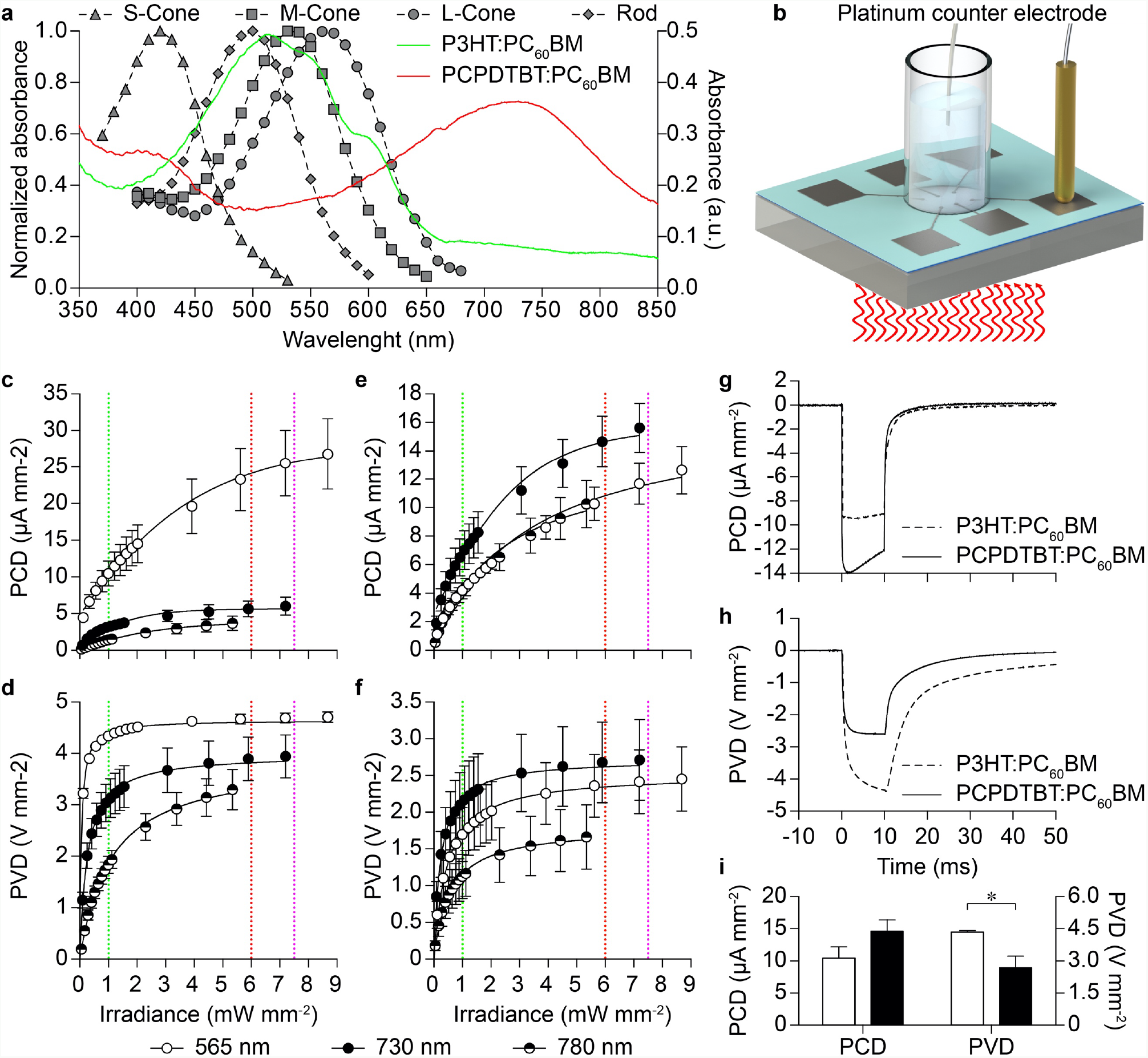
Optimization of the bulk heterojunction. **a**, Normalized absorbance spectra of S-cones, M-cones, L-cones and rods on the left axis (redrawn from ^18^). Absorbance spectra of the P3HT:PC_60_BM and the PCPDTBT:PC_60_BM BHJs on the right axis. **b**, Sketch of the PC and PV recording setup. **c,d**, Mean (± s.e.m., *n* = 4 chips) PCD (**c**) and PVD (**d**) with HTL/P3HT:PC_60_BM/Ti. **e,f**, Mean (± s.e.m., *N* = 3 chips) PCD (**e**) and PVD (**f**) with HTL/PCPDTBT:PC_60_BM/Ti. In panels **c**-**f**, the vertical dotted lines represent the half of the MPE respectively for 565 nm (green, 1 mW mm^−2^), 730 nm (red, 6 mW mm^−2^), and 780 nm (magenta, 7.5 mW mm^−2^). Lines are the interpolations with an asymmetrical, five-parameter, logistic dose-response function for PCDs and a hyperbola function for PVDs. **g,h**, Grand-average PCD (**g**) and PVD (**h**) traces with HTL/P3HT:PC_60_BM/Ti (dashed line, 565 nm, *n* = 4 chips) and HTL/PCPDTBT:PC_60_BM/Ti (solid line, 730 nm, *n* = 3 chips) BHJs at half of the respective MPE. **i**, Mean (± s.e.m.) PCD and PVD obtained with HTL/P3HT:PC_60_BM/Ti (white bars, 565 nm, *N* = 4) and HTL/PCPDTBT:PC_60_BM/Ti (black bars, 730 nm, *n* = 3) at half of the respective MPE.

In this article, we demonstrate that implantable stimulating devices based on conjugated polymers can operate in the NIR range (i.e. wavelength longer than 700 nm). We also document the relevance of the electrical and adhesive properties of the conjugated polymers in the fabrication of an implant with tailored photovoltaic characteristics that achieve an efficient neural stimulation and high stability in aqueous solution. As proof-of-principle, we designed an organic photovoltaic interface for retinal stimulation, based on the poly[2,6-(4,4-bis-(2-ethylhexyl)-4*H*-cyclopenta [2,1-*b*;3,4-*b*′]dithiophene)-*alt*-4,7(2,1,3-benzothiadiazole)] blended with PC_60_BM (PCPDTBT: PC_60_BM) BHJ^1,20^. We demonstrate that this new BHJ can generate electrical responses at safe irradiance levels suitable for retinal stimulation. Furthermore, we investigated the impact of a cross-linking molecule on adhesive properties and morphology of the organic interface. Last, we also verified that this BHJ is not cytotoxic.

## Results

### Optimization of the bulk heterojunction

In order to evaluate the performance of NIR-responsive BHJs, we fabricated chips embedding 6 photovoltaic pixels composed of three layers: an anode made of PEDOT:PSS (HTL Solar), the BHJ (both P3HT:PC_60_BM or PCPDTBT: PC_60_BM), and a cathode made of titanium (Ti). Each photovoltaic pixel was connected to a contact pad to measure its output signal with respect to a platinum counter electrode immersed in saline solution (**Fig. 1b**). We measured the responses of photovoltaic pixels based on both BHJs upon 10-ms light pulses at two NIR wavelengths (730 nm and 780 nm, where the response of cones should be minimal) and compared them to the ones obtained upon green light stimulation (565 nm). In all the experiments, for each chip, the responses from the 6 electrodes were measured, the peak amplitudes were quantified, and the data were averaged.

As expected, the P3HT:PC_60_BM BHJ showed the strongest photo-current density (PCD) at 565 nm (**Fig 1c**, white circles). However, to evaluate the three wavelengths in perspective of retinal prosthesis needs, we compared the PCD values obtained at the irradiance corresponding to half of the respective MPE (1, 6, and 7.5 mW mm^−2^ respectively for 565, 730, and 780 nm). The mean PCD obtained at 565 nm with pulses of 1 mW mm^−2^ (white circles, green line) is 1.85 times higher than the mean PCD obtained at 730 nm (black circles, red line) with pulses of 6 mW mm^−2^. At 780 nm (black/white circles) the maximal irradiance obtained from the LED was 5.35 mW mm^−2^, which is lower than the half of the MPE (7.5 mW mm^−2^). Therefore, the theoretical PCD value was computed using the interpolating curve (R^2^ = 0.74). In both cases (730 and 780 nm), the PCD obtained at half of the MPE is largely lower than the one obtained with green light (565 nm). The photo-voltage density (PVD) had a similar behavior (**Fig. 1d**). The theoretical PVD at 780 nm and 7.5 mW mm^−2^ (black/white circles, magenta line) was computed using the interpolating curve (R^2^ = 0.90). Based on these results, the use of the HTL/P3HT:PC_60_BM/Ti pixels appears not ideal for a NIR-responsive retinal prosthesis. Hence, we investigated a different BHJ based on the PCPDTBT:PC_60_BM blend.

In this case, the wavelength with the highest PCD (**Fig. 1e**) and PVD (**Fig. 1f**) is 730 nm (black circles). Moreover, if compared at half of the MPE, the PCD obtained at 730 nm with HTL/PCPDTBT:PC_60_BM/Ti is 1.4 times higher than the PCD obtained at 565 nm with HTL/P3HT:PC_60_BM/Ti (**Fig. 1g,i**; two-tailed t-test, p = 0.1571). On the other hand, the PVD is slightly lower (**Fig. 1h,i**; two-tailed t-test, p < 0.05). Despite these very small differences, this suggests that the PCPDTBT:PC_60_BM BHJ should be as efficient in retinal stimulation as the traditional P3HT:PC_60_BM blend, considering adjusted irradiances to their respective MPEs. In addition, as previously reported for HTL/P3HT:PC_60_BM/Ti pixels, the photovoltaic electrodes based on the PCPDTBT:PC_60_BM BHJ are fully discharged in less than 40 ms from the pulse offset (**Fig. 1h**); this is an important feature allowing a stimulation pulse rate up to 20 Hz, which is ideal for retinal prosthesis ^12^.

### Optimization of the anodic layer

Photovoltaic organic retinal prostheses typically rely on a bottom anode made of PEDOT:PSS ^5,12–14,16^. We therefore investigated the effect of its conductivity by using two formulations from Clevios Heraeus: HTL Solar (0.47 S mm^−1^, average of 5 measures from 1 sample; film thickness of 60 nm) and PH1000 (1.17 S mm^−1^, average of 5 measures from 1 sample; film thickness of 90 nm). At 730 nm, the mean PCD obtained with the PH1000/PCPDTBT:PC_60_BM/Ti pixel (grey squares) is considerably higher than the HTL/PCPDTBT:PC_60_BM/Ti pixel (black circles), while the mean PVD increases with a lower rate but it reaches the same value at half of the MPE for 730 nm (**Fig. 2a-c**). To summarize our findings (**Fig. 2d,e**), the PVD and PCD values obtained with the three configurations presented were compared at their best operational wavelengths (565 nm for P3HT and 730 nm for PCPDTBT) and at half of the respective MPE (1 mW mm^−2^ for 565 nm and 6 mW mm^−2^ for 730 nm). The PH1000/PCPDTBT:PC_60_BM/Ti pixels (grey squares) showed the highest PCD (one-way ANOVA, p = 0.0018; Tuckey’s multiple comparisons test: HTL/P3HT:PC_60_BM/Ti *vs* HTL/PCPDTBT:PC_60_BM/Ti p = 0.9235, HTL/P3HT:PC_60_BM/Ti *vs* PH1000/PCPDTBT:PC_60_BM/Ti p = 0.0024, HTL/PCPDTBT:PC_60_BM/Ti *vs* PH1000/PCPDTBT:PC_60_BM/Ti p = 0.0062). On the other hand, the HTL/P3HT:PC_60_BM/Ti pixels (white circles) showed the highest PVD (one-way ANOVA, p = 0.0034; Tuckey’s multiple comparisons test: HTL/P3HT:PC_60_BM/Ti *vs* HTL/PCPDTBT:PC_60_BM/Ti p = 0.0105, HTL/P3HT:PC_60_BM/Ti *vs* PH1000/PCPDTBT:PC_60_BM/Ti p = 0.0046, HTL/PCPDTBT:PC_60_BM/Ti *vs* PH1000/PCPDTBT:PC_60_BM/Ti p = 0.9567). The HTL/PCPDTBT:PC_60_BM/Ti pixels (black circles) showed the lowest PCD and PVD. Interestingly, the different configurations also have different photovoltage discharge rates at the offset of the light pulse (**Fig. 2e**). The PH1000/PCPDTBT:PC_60_BM/Ti pixel showed the faster discharge rate (two phase exponential decay, τ_fast_ = 0.85 and τ_slow_ = 3.99, R^2^ = 0.8), probably because of the higher electrical conductivity of PH1000.

**Figure 2.**
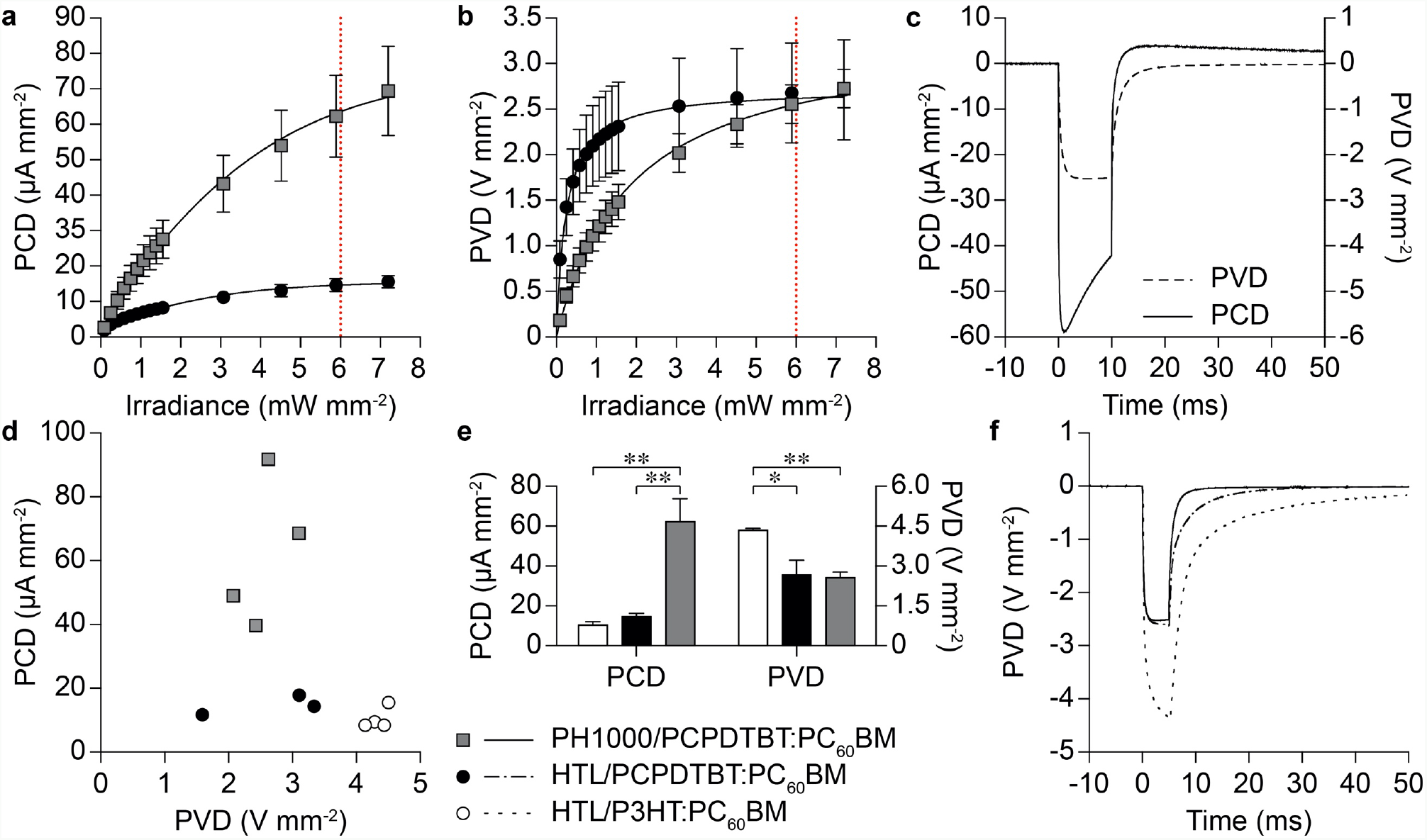
Optimization of the anodic layer. **a,b**, Mean (± s.e.m.) PCD (**a**) and PVD (**b**) obtained at 730 nm for HTL/PCPDTBT:PC_60_BM/Ti (black circles, *n* = 3 chips) and PH1000/PCPDTBT:PC_60_BM/Ti (grey squares, *n* = 4 chips). The solid lines are the interpolations with an asymmetrical, five-parameter, logistic dose-response function for PCDs and a hyperbola function for PVDs. The vertical red dotted lines represent the half of the MPE for 730 nm (6 mW mm^−2^). **c**, Grand-average PCD (solid line) and PVD (dashed line) traces obtained with PH1000/PCPDTBT:PC_60_BM/Ti with 730 nm at half of the MPE (*n* = 4 chips). **d**, PVD/PCD plot for HTL/P3HT:PC_60_BM/Ti (white circles, *n* = 4 chips, 565 nm, 1 mW mm^−2^), HTL/PCPDTBT:PC_60_BM/Ti (black circles, *n* = 3 chips, 730 nm, 6 mW mm^−2^), and PH1000/PCPDTBT:PC_60_BM/Ti (grey squares, *n* = 4 chips, 730 nm, 6 mW mm^−2^). **e**, Mean (± s.e.m.) PCD and PVD obtained with HTL/P3HT:PC_60_BM/Ti (white bars, 10.46 ± 1.72 μA mm^−2^, 4.34 ± 0.08 V mm^−2^, *n* = 4 chips, 565 nm, 1 mW mm^−2^), HTL/PCPDTBT:PC_60_BM/Ti (black bars, 14.65 ± 1.79 μA mm^−2^, 2.68 ± 0.55 V mm^−2^, *N* = 3 chips, 730 nm, 6 mW mm^−2^), and PH1000/PCPDTBT:PC_60_BM/Ti (grey bars, 62.28 ± 11.54 μA mm^−2^, 2.56 ± 0.22 V mm^−2^, *n* = 4 chips, 730 nm, 6 mW mm^−2^) at half of the MPE respectively for 565 nm and 730 nm. **f**, Comparison of the PVD grand-average traces for HTL/P3HT:PC_60_BM/Ti (dotted line, *n* = 4 chips, 565 nm, 1 mW mm^−2^), HTL/PCPDTBT:PC_60_BM/Ti (dashed-dotted line, *n* = 3 chips, 730 nm, 6 mW mm^−2^), and PH1000/PCPDTBT:PC_60_BM/Ti (solid line, *n* = 4 chips, 730 nm, 6 mW mm^−2^).

### Optimization of the adhesion and stability of the interface

Strong adhesion and stability are a prerequisite for the long-term functioning of an implantable device. However, in organic-based prostheses, the adhesion and stability of the PEDOT:PSS layer over a substrate in aqueous environment is limited by the delamination and solubility of PSS. A common strategy to obtain water-stable thin films of PEDOT:PSS is to add the (3-glycidyloxypropyl)trimethoxysilane (GOPS) crosslinker (typically 1 v/v%), which prevents both dissolution and delamination of PEDOT:PSS films^21^. On the other hand, it was also reported that the electrical conductivity of PEDOT:PSS films decreases as a function of the GOPS content^21,22^. We found that the addition of 1 v/v% of GOPS to PH1000 has an impact on the PCD (**Fig. 3a,c**) and PVD (**Fig. 3b,d**) generated by the photovoltaic electrodes. In particular, for the PH1000/PCPDTBT:PC_60_BM/Ti pixels, the addition of 1 v/v% of GOPS (grey circles) reduces the PCD and increases the PVD at 730 nm. Moreover, it causes a very slow photovoltage discharge (two phase exponential decay, τ_fast_ = 11.35 and τ_slow_ = 77.38, R^2^ = 0.98), which does not allow the generation of distinct voltage pulses at 20 Hz of repetition rate (**Fig. 3e**). Indeed, such configuration does not allow a full discharge and recharge of the electrode voltage between each pulse during 20 Hz train stimulation, in contrast to the case when GOPS is not added (**Fig. 3f**). This slower discharge rate is very likely caused by the reduction of the electrical conductivity due to the addition of 1 v/v% of GOPS (0.21 S mm^−1^, average of 5 measures from 1 sample; film thickness of 170 nm).

**Figure 3.**
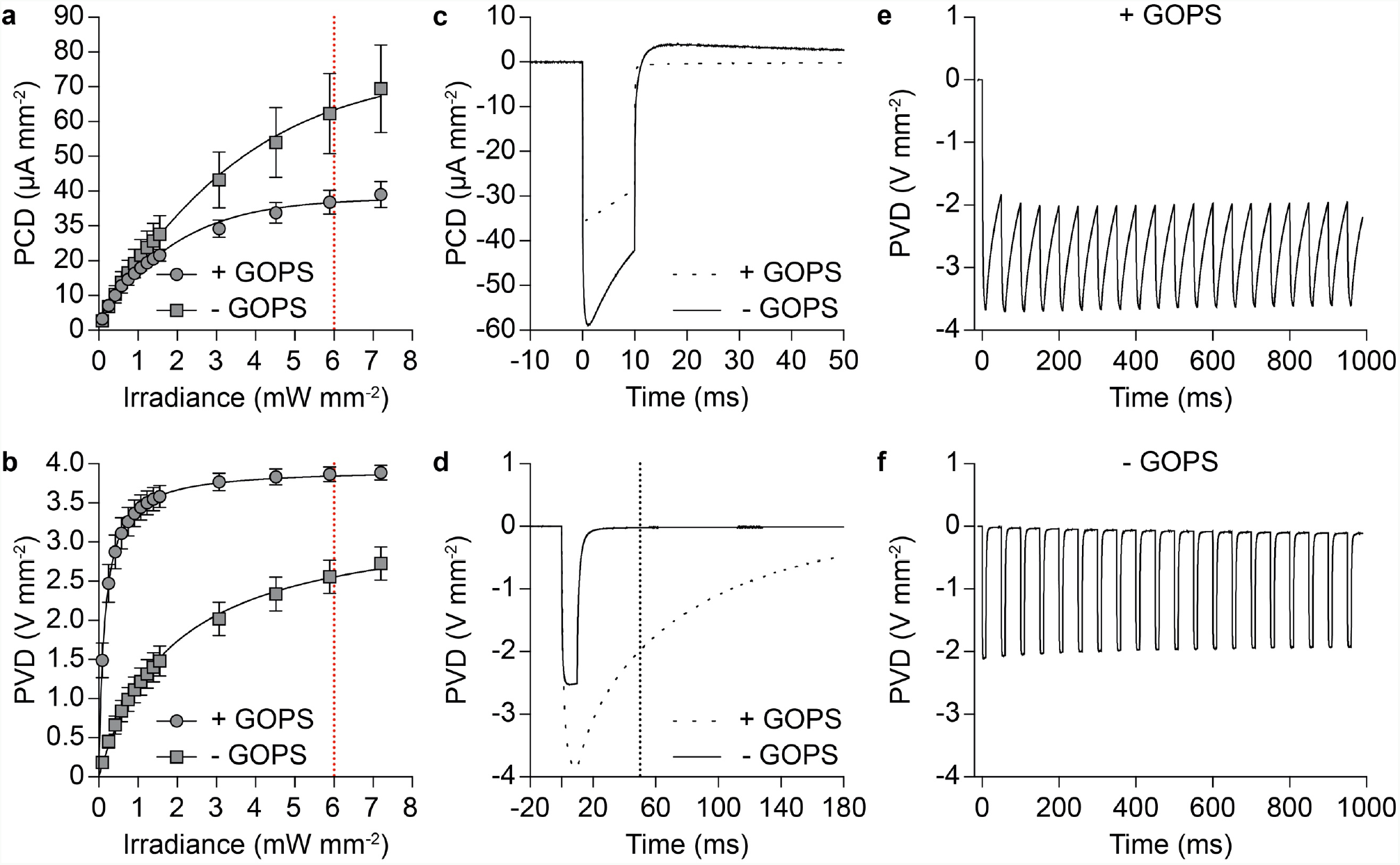
Optoelectronic responses with GOPS. **a,b**, Mean (± s.e.m.) PCD (**a**) and PVD (**b**) obtained with pristine PH1000/PCPDTBT:PC_60_BM/Ti (grey squares, *n* = 4) and PH1000/PCPDTBT:PC_60_BM/Ti supplemented with 1 v/v% GOPS (grey circles, *n* = 3) at 730 nm. The solid lines are the interpolations with an asymmetrical, five-parameter, logistic dose-response function for PCDs and a hyperbola function for PVDs. The vertical red dotted lines represent the half of the MPE for 730 nm (6 mW mm^−2^). **c,d**, Grand-average PCD (**c**) and PVD (**d**) traces obtained upon 10-ms pulses at 730 nm and half of the MPE (6 mW mm^−2^) with pristine PH1000 (solid line, *n* = 4) and PH1000 with GOPS (dashed line, *n* = 3). **e,f**, Representative traces from single electrodes of PVD upon the delivery of a train stimulation composed of 20 pulses (10 ms, 730 nm, 6 mW mm^−2^) delivered at 20 Hz with pristine PH1000 (**f**) and PH1000 with 1 v/v% of GOPS (**e**).

Therefore, we investigated which concentration of GOPS could simultaneously increase the stability and adhesion of the PEDOT:PSS film (PH1000) while preserving a photovoltaic performance suitable for retinal stimulation. We fabricated a set of samples with five concentration of GOPS (0, 0.1, 0.25, 0.5, and 1 v/v%) and measured the PCD and PVD. The mean PVD (730 nm, 6 mW mm^−2^) increased immediately upon addition of GOPS and it remained stable regardless of the concentration (**Fig. 4a**). Conversely, the mean PCD peak remain high up to 0.1 v/v% of GOPS and then it decreased. Therefore, one can designate a concentration of 0.1 v/v% as the best compromise, since higher concentration would induce a strong reduction of the PCD generated by the photovoltaic pixels. In parallel, the increase in the concentration of GOPS increases the decay time at the offset of the light pulse (**Fig. 4b**). The fitting with a two-phase exponential decay function showed that both τ_fast_ and τ_slow_ increase with the GOPS concentration (**Fig. 4c**). However, with 0.1 v/v% of GOPS, the electrode is fully discharged (i.e., the PV returns to baseline) in 40 ms from the pulse offset. This condition represents the optimal situation for a retinal implant because the fast photovoltage decay allows the precise reproduction of 20-Hz pulse trains (**Fig. 4d**). In summary, the chemical modification of PEDOT:PSS with 0.1 v/v% of GOPS allowed the tailoring of the photovoltaic performance for an efficient retinal stimulation. Both the resistance (**Fig. 4e**, white circles) and the thickness (**Fig. 4f**) of the PEDOT:PSS thin film are affected by the GOPS concentration. Accordingly, the film conductivity (computed by normalizing the average film resistance by the average film thickness) decreases with the increase of the GOPS concentration (**Fig. 4e**, black circles). This confirms that the conductivity of the PEDOT:PSS films plays a major role in the optoelectronic properties of photovoltaic pixels.

**Figure 4.**
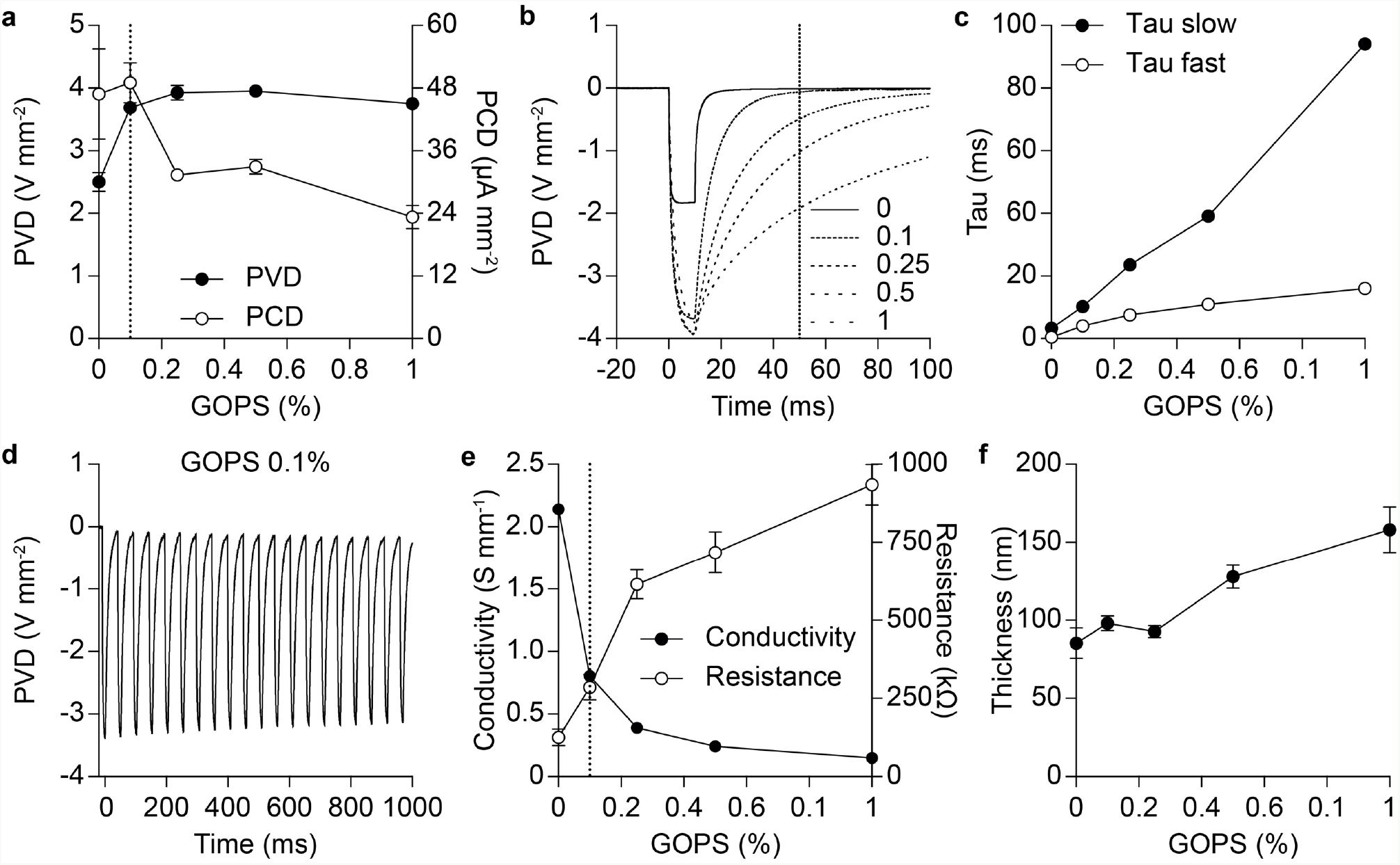
Electrical tuning with the GOPS concentration. **a**, Mean (± s.e.m.) PCD (right axis) and PVD (left axis) obtained with PH1000/PCPDTBT:PC_60_BM/Ti supplemented with various concentrations of GOPS (0, 0.1, 0.25, 0.5, 1 v/v%, *n* = 4 chips for each concentration) at 730 nm and half of the MPE (6 mW mm^−2^). **b**, Grand-average (*n* = 4 chips for each concentration) PVD traces obtained at the same conditions as in (**a**). **c**, Evolution of τ_fast_ and τ_slow_ as a function of the GOPS concentration. **d**, Representative traces from a single electrode of PVD upon 20 pulses of 10 ms at 730 nm at half of the MPE (6 mW mm^−2^) and delivered at 20 Hz with PH1000 supplemented with 0.1 v/v% of GOPS. **e**, Mean (± s.d.; 12 measures from *n* = 2 samples for each condition) resistance (white circles, right axis) and average conductivity (black circles, left axis) of PH1000 films with various concentrations of GOPS. **f**, Mean (± s.d.; 12 measures from *n* = 2 samples for each condition) thickness of PEDOT:PSS (PH1000) films with various concentrations of GOPS.

To further investigate the role of GOPS in the adhesion and stability of the PEDOT:PSS layer, we fabricated full POLYRETINA devices with the PCPDTBT:PC_60_BM but without encapsulation of the active layer (**Fig. 5a,b**)^12^. The lack of polydimethylsiloxane (PDMS) encapsulation allows a faster delamination due to the direct contact of the organic layers with water. Soaking experiments in saline solution revealed that the addition of 0.1 v/v% of GOPS (**Fig. 5c**, middle) increases the stability of the photovoltaic pixels compared to pristine PEDOT:PSS (**Fig. 5c**, left). On the other hand, a higher concentration of GOPS (0.25 v/v%; **Fig. 5c**, right) does not increase adhesion but induces again delamination. In fact, the addition of GOPS not only crosslinks the PSS molecules together, but it also anchors them to the substrate (e.g. PDMS). This explains the strengthened adhesion of PEDOT:PSS to PDMS. Nevertheless, a further increase in the GOPS concentration (e.g. from 0.1 to 0.25 v/v% and above) induced delamination again, but at the interface between PEDOT:PSS and PCPDTBT:PC_60_BM rather than at the interface with PDMS.

**Figure 5.**
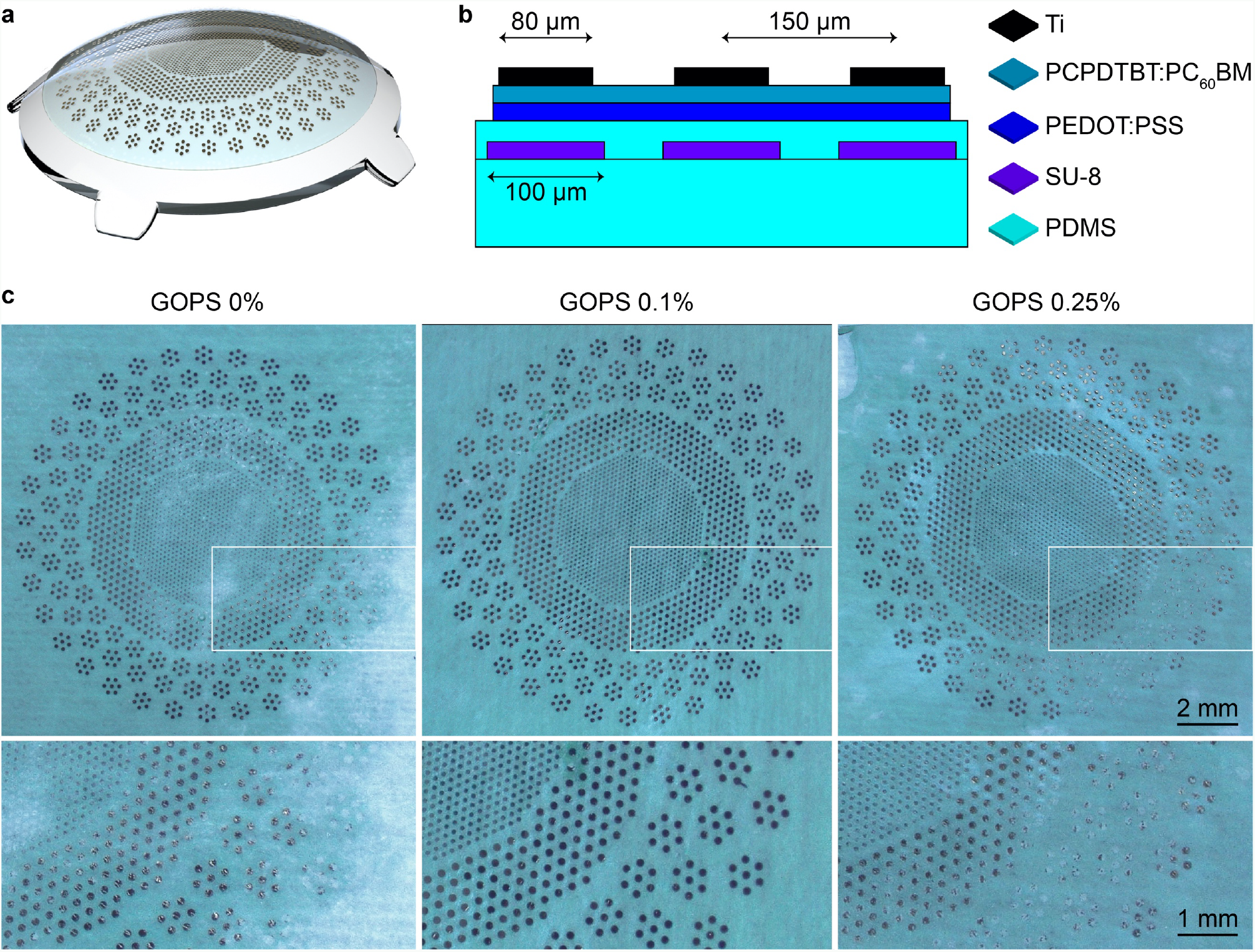
Adhesion of the interface. **a**, 3D model of the POLYRETINA prosthesis. **b**, Cross-section of the POLYRETINA active interface, including: PDMS (50 μm), a second layer of PDMS (14 μm) embedding SU-8 rigid platforms (6 μm), a layer of PEDOT:PSS with various percentages of GOPS, a layer of PCPDTBT:PC_60_BM (100 nm), and titanium cathodes (150 nm). **c**, Pictures of POLYRETINA devices prepared with 0, 0.1, and 0.25 v/v% of GOPS after soaking in saline solution and ultra-sonication for 5 min.

We hypothesize that the increase of the cross-linking degree with GOPS could impair the diffusion of the PCPDTBT molecules into PEDOT:PSS during the thermal treatment after deposition. This lower interaction between the two polymeric layers would inevitably decreases the adhesion forces. To verify this hypothesis, we performed depth-profiling measurements with time-of-flight secondary ion mass spectrometry (ToF-SIMS) from the top surface of PCPDTBT:PC_60_BM to the bottom side of PEDOT:PSS (i.e., to the substrate) at various concentration of GOPS (**Fig. 6a**). The analysis of the negative polarity spectra indicates that with increasing GOPS concentration, PC_60_BM molecules (red curve, C_60_^−^ fragments) tend to accumulate at the interface towards PEDOT:PSS. Furthermore, the depth distribution of the CN^−^ fragments (green curve) suggests a penetration of PCPDTBT molecules into the PEDOT:PSS layer when GOPS is not added. Based on the depth profiles of the relevant fragments, we predicted the organization of the organic molecules and represent it for the two extreme GOPS concentrations: 0 and 1 v/v% (**Fig. 6b**). In **Fig. 6c**, the normalized intensities of the PCPDTBT signal (CN^−^ fragments) for the three different GOPS concentrations are plotted to efficiently compare their slopes, representing their distinct penetration depths into PEDOT:PSS. Moreover, the slope of the normalized intensities of the PEDOT signal (SC_2_O^−^ fragments) also shows a diffuse interface (slow rise) for 0 v/v% and a sharper one (faster rise) for 1 v/v% (**Fig. 6d**). We therefore assume that PCPDTBT molecules are more easily able to penetrate the PEDOT:PSS network if the latter is less cross-linked, allowing a stable interface against delamination. Conversely, if PEDOT:PSS is supplemented with GOPS, PC_60_BM is placed at the interface with PEDOT:PSS, creating a brittle and more fragile interface where delamination can occur ^23^. Hence, a concentration of 0.1 v/v% of GOPS is optimal to enhance both the photovoltaic performance for retinal stimulation and the stability of the two interfaces.

**Figure 6.**
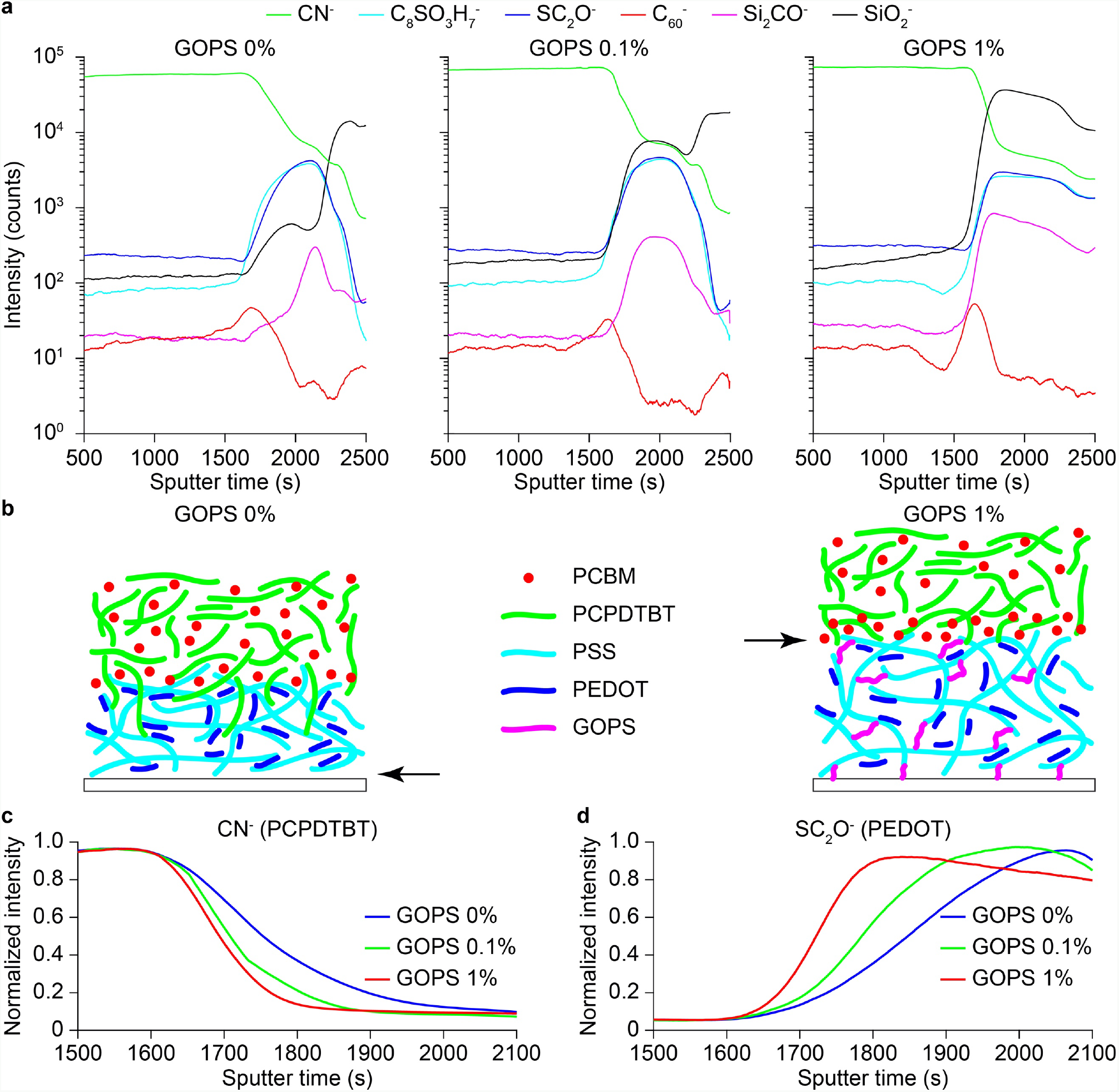
Redistribution of molecules as a function of the GOPS concentration. **a**, Smoothed ToF-SIMS depth profiles (negative polarity) conducted on the organic layers with 0, 0.1, and 1 v/v% of GOPS. CN^−^, C_8_SO_3_H_7_^−^, SC_2_O^−^, C_60_^−^, Si_2_CO^−^, and SiO_2_^−^ are used as proxies respectively for PCPDTBT, PSS, PEDOT, PC_60_BM, GOPS, and both glass and GOPS. The interface between PCPDTBT: PC_60_BM and PEDOT:PSS is roughly at 1700 s, while the one between PEDOT:PSS and the glass substrate is roughly at 2100 s, 2250 s, and 2400 s for pristine PEDOT:PSS, with 0.1 v/v% GOPS, and with 1 v/v% GOPS, respectively. **b**, Schematic representation of the deduced organization of the organic molecules for 0 v/v% (left) and 1 v/v% (right) of GOPS. The arrows indicate the observed delamination level within the organic layers. **c,d**, Normalized intensities of the depth profiles for PCPDTBT (CN^−^) (**c**) and PEDOT (SC_2_O^−^) (**d**) for the three GOPS concentrations.

### Functional validation of the near-infrared-responsive photovoltaic prosthesis

PCPDTBT was used for bioelectronic interfaces only in few reports ^1,10^. Therefore, we first investigated the cytotoxicity of the near-infrared-responsive photovoltaic prosthesis fabricated above a PDMS substrate (**Fig. 7a**). A mean cell viability (± s.e.m.; *N* = 4 samples) of 107.725 ± 0.520 was obtained (negative control 100 %, *N* = 1 sample; positive control 0 %, *N* = 1 sample), thus confirming the non-toxicity of the prosthesis. For each sample the test was performed on triplicate cultures wells and data were averaged (**Fig. 7b**). A one-way ANOVA analysis (p < 0.0001, F = 223.9) revealed that all the four prostheses tested resulted in a cell viability significantly higher than the positive control (p < 0.0001 for all, Tukey multiple comparisons); similarly, the negative control is significantly higher than the positive control (p < 0.0001, Tukey multiple comparisons). There is no statistically significant difference among the four prostheses and against the negative control (1 vs 2: p = 0.9947; 1 vs 3: p > 0.9999; 1 vs 4: p > 0.9999; 2 vs 3: p = 9989; 2 vs 4: p = 0.9920; 3 vs 4: p > 0.9999; 1 vs negative control: p = 0.5401; 2 vs negative control: p = 0.2888; 3 vs negative control: p = 0.4598; 4 vs negative control: p = 0.5678; Tukey multiple comparisons).

**Figure 7.**
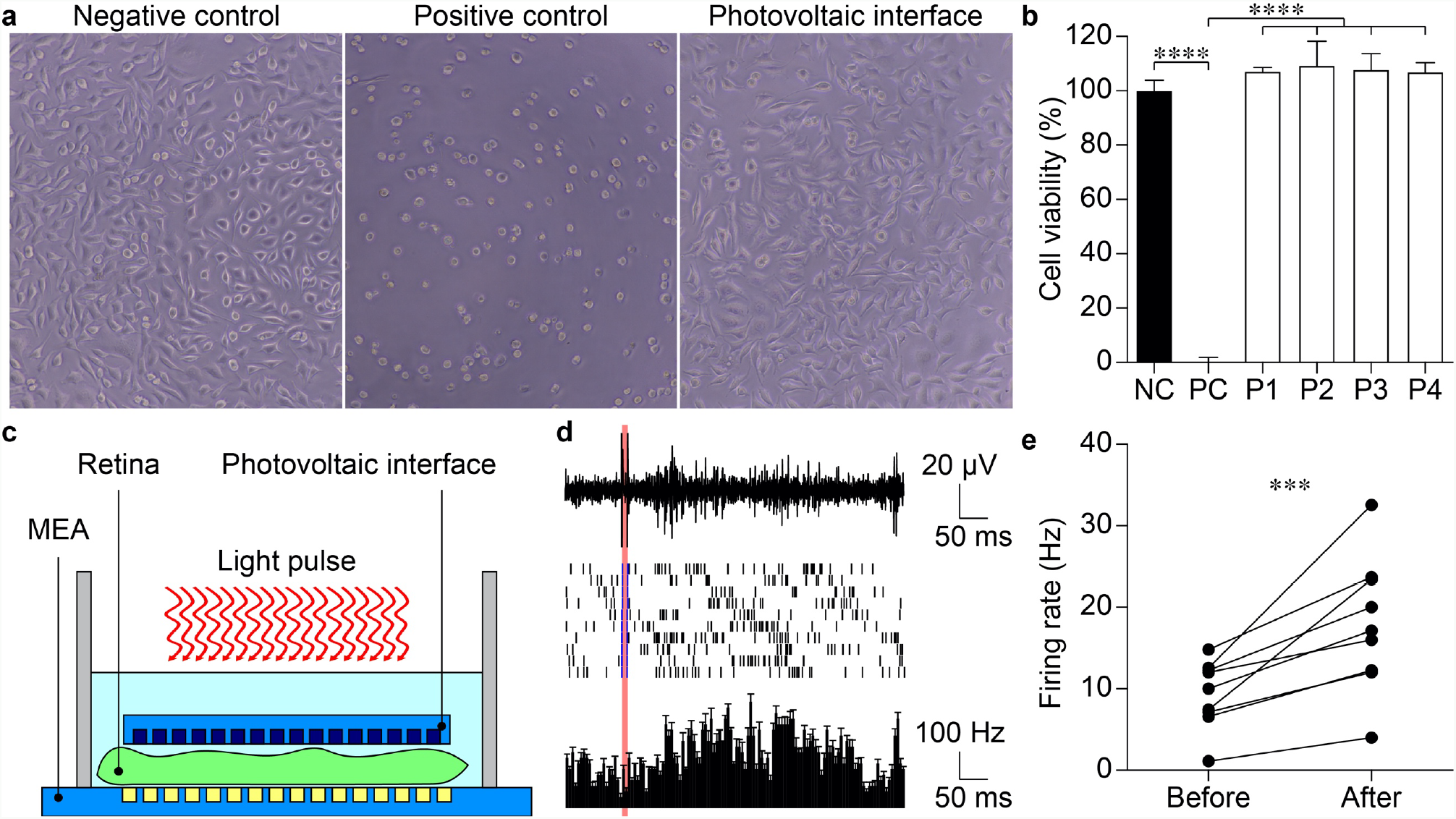
Functional validation *in-vitro* and *ex-vivo*. **a**, Representative images of *in-vitro* cytotoxicity of the near-infrared-responsive photovoltaic prosthesis (right). Optical images of cell culture as negative control and positive controls are also shown (left and middle respectively). **b**, Quantification of cell viability of the four prostheses tested (from P1 to P4) along with the negative and positive controls (respectively NC and PC). **c**, Sketch of the setup used for *ex-vivo* electrophysiology. **d**, Representative response of one retinal ganglion cell upon NIR stimulation. The top row shows the electrophysiological recordings upon one light pulse (10 ms, 4.7 mW mm^−2^; identified by the red bar). The middle row shows the raster plots upon ten consecutive NIR light pulses. The blue lines in the raster correspond to the events classified as stimulus artefacts. The bottom row shows the corresponding mean post-stimulus time histogram (± s.d., bins of 5 ms). **e**, Quantification of the network-mediated activity in retinal ganglion cells, before and after the photovoltaic stimulation (*n* = 9 retinal ganglion cells; one-tailed paired t-test, p < 0.001).

Last we verified that the near-infrared-responsive photovoltaic prosthesis was able to efficiently stimulate retinal cells. Explanted retinas from 4-months old retinal degeneration 10 mice, an established model for retinitis pigmentosa ^24–26^, were placed onto a microelectrode array (MEA) and the responses evoked by photovoltaic stimulation (10 ms light pulses, 4.7 mW mm^−2^) were recorded from retinal ganglion cells (**Fig. 7c**). We previously demonstrated that at this age, retinas have no light responsivity ^27^. NIR light pulses were able to significantly induce network mediated firing activity in retinal ganglion cells (**Fig. 7d,e**) at a safe irradiance level (2.5 times lower than the MPE), thus showing that the near-infrared-responsive photovoltaic prosthesis can effectively be used for retinal stimulation.

## Discussion

Photovoltaics is an attractive approach in bioelectronic medicine and neuroprosthetics to stimulate or modulate neuronal activity. Our results show that organic photovoltaic interfaces can be tailored to achieve higher stability, better optoelectronic performances, and adjusted sensitivity in order to match the desired target application.

In a proof-of-concept, we demonstrated the advantage of a near-infrared-responsive neuroprosthesis for retinal stimulation that allows for higher compliance with the standard for optical safety and avoids the interfering with the residual natural vision. So far, inorganic photovoltaic retinal prostheses were considered better suited for photovoltaic stimulation because of the higher NIR sensitivity of silicon ^28^. Previous researches attempted to perform retinal stimulation at longer wavelengths, even if still in the visible range ^29^, and a computational study showed that a photovoltaic interface based on conjugated polymers could operate in the NIR range ^30^; however, in this study an experimental validation was not provided. We demonstrated that also organic photovoltaic interfaces can efficiently stimulate blind retinas at NIR wavelengths (i.e. 730 nm). This represents an important contribution to the development of organic retinal implants. Organic photovoltaic bioelectronic interfaces are facing additional challenges, such as tailoring the electrical properties of the photovoltaic cell to meet the required working conditions and the weak stability of the organic materials due to water-induced swelling, degradation, and delamination. Our study addressed those open challenges to advance the exploitation of conjugated polymers in photovoltaic organic prostheses.

## Acknowledgments

We would like to acknowledge the EPFL Center of Micronanotechnology and the The Neural Microsystems Platform at Wyss Center for their support. We acknowledge also Prof. Kevin Sivula and Dr. Nestor Guijarro Carratala of EPFL for the support with the UV-Vis spectroscopy measurements. This work has been supported by École polytechnique fédérale de Lausanne, Medtronic, European Commission (Project 701632), Fondation Mercier pour la science, Velux Stiftung (Project 1102), Gebert Rüf Stiftung (Project GRS-035/17), and Swiss National Science Foundation (Project CR23I2-162828).

## Author contributions

M.J.I.A.L. fabricated the samples and devices, recorded the photo-voltage and the photo-current, measured the thickness and resistance, and performed the soaking experiments. N.A.L.C. performed *ex-vivo* electrophysiology. L.F. fabricated the samples and performed the absorbance measurements. M.K. performed time-of-flight secondary ion mass spectrometry measurements. E.G.Z. performed *in-vitro* cytotoxicity. D.G. supervised the entire study and wrote the manuscript. All the authors read and accepted the manuscript.

## Data and materials availability

All data needed to evaluate the conclusions in the paper are present in the paper. Additional data related to this paper may be requested from the corresponding author upon reasonable request.

## Materials and Methods

### Chip micro-fabrication

Samples were fabricated on 20 × 24 mm^2^ glass substrates (2947-75X50, Corning Incorporated) cleaned by ultra-sonication in acetone, isopropyl alcohol, and deionized water for 15 min each and then dried with nitrogen gun. The deposition of the PEDOT:PSS and the preparation of the bulk heterojunctions were performed in a glovebox under nitrogen atmosphere. PEDOT:PSS (HTL Solar and PH1000, Clevios) was filtered (1 μm PTFE filters) then spin-coated at 3000 rpm for 40 seconds on each sample. Subsequent annealing at 115 °C for 30 min was performed. When present, GOPS was added to the solution before filtering. 20 mg of P3HT (698997, Sigma Aldrich) or PCPDTBT (754005-100MG, Sigma) and 20 mg of PC_60_BM (M111, Ossila) were dissolved in 1 mL of anhydrous chlorobenzene each and let stirring overnight at 70 °C. The solutions were then filtered (0.45 μm PTFE filters) and blended [1:1 v:v]. The P3HT:PC_60_BM (nominal thickness 100 nm) and PCPDTBT:PC_60_BM (nominal thickness 100 nm) blends were then spin-coated at 1000 rpm for 45 seconds. Subsequent annealing at 115 °C for 30 min was performed. Titanium cathodes (diameter 100 μm, nominal thickness 150 nm) were deposited by DC magnetron sputtering through a shadow mask. A plastic reservoir was then attached to the sample using PDMS as adhesive.

### Measure of photo-voltage and photo-current

Samples were placed on a holder, and each electrode was sequentially contacted. A platinum wire immersed in physiological saline solution (NaCl 0.9 %) was used as counter electrode. 10-ms light pulses were delivered by a 565-nm (M565L3, Thorlabs), 730-nm (M730L4, Thorlabs), or 780-nm (M780LP1, Thorlabs) LED focused at the sample level. Photo-voltage and photo-current were measured using respectively a voltage amplifier (1201, band DC-3000 Hz, DL-Instruments) and a current amplifier (1212, DL-Instruments). Data sampling (40 kHz) and instrument synchronization were obtained via a DAQ board (PCIe-6321, National Instruments) and a custom-made software. Data analysis was performed in Matlab (Mathworks). When evaluating the photo-voltage and photo-current densities generated by the interface, also the area of the connecting line exposed to light has been considered.

### Spectral absorbance

The preparation of the bulk heterojunctions was performed as before. The thickness were 80 and 62 nm for P3HT:PC_60_BM and PCPDTBT:PC_60_BM respectively. The absorbance spectra of the thin films were measured using a UV-vis-NIR UV-3600 Shimadzu spectrophotometer.

### Resistance measurements

The preparation of the PEDOT:PSS was performed as before. The film resistance was measured with a custom 4-point prober (2.5 mm pitch distance) using a Keithley 2400 source-meter. Each sample was measured on five different locations randomly distributed on the surface.

### Thickness measurements

Thin film thickness was measured in PeakForce tapping mode (ScanAsyst Air silicon tip, f_0_ = 70 kHz, k = 0.4 N m^−1^) with a Dimension Icon AFM (Bruker).

### Time-of-flight secondary ion mass spectrometry measurements

The measurements were performed on a ToF-SIMS.5 instrument (IONTOF, Germany) operated in the spectral mode using a 25 keV Bi^3+^ primary ion beam with an ion current of 0.81 pA. A mass resolving power in the range of 5000 m Δm^−1^ was reached. For depth profiling, a 500 eV Cs^+^ sputter beam with a current of 43.47 nA was used. The raster area of the sputter beam was 500 μm × 500 μm, and the mass-spectrometry was performed on an area of 200 μm × 200 μm in the center of the sputter crater. A low-energy electron flood gun was used for charge compensation.

### Fabrication of POLYRETINA prostheses

Prostheses were prepared as previously described ^12^. A thin sacrificial layer of poly(4-styrenesulfonic acid) solution (561223, Sigma-Aldrich) was spin-coated on 4” Si wafers (1000 rpm, 40 s) and baked (120 °C, 15 min). Degassed PDMS pre-polymer (10:1 ratio base-to-curing agent, Sylgard 184, Dow-Corning) was then spin-coated (1000 rpm, 60 s) and cured in oven (80 °C, 2 hr). After surface treatment with oxygen plasma (30 W, 30 s), a 6 μm thick SU-8 (GM1060, Gersteltec) layer was spin-coated (3800 rpm, 45 s), soft-baked (110 °C, 300 s), exposed (140 mJ cm^−2^, 365 nm), post-baked (90 °C, 1800 s; 60 °C, 2700 s), developed in propylene glycol monomethyl ether acetate (48443, Sigma-Aldrich) for 2 min, rinsed in isopropyl alcohol, and dried with nitrogen gun. After surface treatment with oxygen plasma (30 W, 30 s), a second layer of degassed PDMS pre-polymer (10:1) was spin-coated (3700 rpm, 60 s) and cured in oven (80 °C, 2 hr). The PEDOT:PSS (PH1000) film, the PCPDTBT:PC_60_BM film, and the titanium cathodes were prepared as described above. The photovoltaic membrane was then released from the wafer and plasma bonded to a PDMS dome-shaped support with a 12 mm curvature radius.

### Soaking experiments

POLYRETINA devices were soaked in saline solution (NaCl 0.9 %) at 37 °C and ultra-sonication was applied for 5 min.

### Cytotoxicity test

The test was conducted according to the requirement of ISO 10993-5: Biological Evaluation of Medical Devices, *in-vitro* cytotoxicity test. Prostheses were sterilized in a dry oven for 2 hr at 120 °C. The test on extraction was performed with samples for a total surface area of 3.54 cm^2^ each, with a ratio of the product to extraction vehicle of 3 cm^2^ ml^−1^. Extraction vehicle was Eagle's Minimum Essential Medium supplemented with fetal bovine serum, penicillin-streptomycin, amphotericin B, and L-glutamine. The extraction was performed for 24 hr at 37 °C. For each sample, the extract was added on triplicate cultures wells containing a sub-confluent L929 cell monolayer. The test samples and the control wells were incubated at 37 °C in 5 % CO_2_ for 24 hours. Following incubation, the cell cultures were examined for quantitative cytotoxic evaluation. 50 μl per well of XTT reagent were added to the cells then incubated at 37 °C in 5 % CO_2_ for further 3 to 5 hr. An aliquot of 100 μl was then transferred from each well into the corresponding wells of a new plate and the optical density was measured at 450 nm.

### Electrophysiology ex-vivo

Animal experiments were performed according to the animal authorizations GE3717 issued by the Département de l’Emploi, des Affaires sociales et de la Santé (DEAS), Direction Générale de la Santé of the Republique et Canton de Genève (Switzerland). Male and female mice from a homozygous colony of retinal degeneration 10 mice (B6.CXB1-Pde6b^rd10^/J, The Jackson Laboratory, Stock number: 004297) were used for the experiments. All animals were kept in a 12 h day/night cycle with access to food and water *ad libitum*. All the experiments were carried out during the day cycle. Eyes were enucleated from euthanized mice (sodium pentobarbital, 150 mg kg^−1^) and dissected in carboxygenated (95% O_2_ and 5% CO_2_) Ames’ medium (A1420, Sigma-Aldrich) under dim red light. Retinas were placed ganglion cells down and maintained in contact with a transparent microelectrode array with 256 electrodes (256MEA200/30iR-ITO, Multi Channel Systems). The near infra-red responsive, polymeric and photovoltaic neuroprosthesis was placed on top of the retina, and both the retina and the prosthesis were kept in position using a 1 mm nylon mesh. Retinas were continuously superfused with carboxygenated Ames’ medium at 32 °C and maintained under dim red light during all the experiments. Light stimuli were generated using a 730-nm light emitting diode (M730L4, Thorlabs) paired to a 20x objective (CFI Plan Apochromat Lambda, Nikon Instruments). The diameter of the light spot was 4.16 mm. The signal from the 256 recording electrodes was amplified, filtered (300 – 3000 Hz), and digitalized at 10 kHz (USB-MEA256-System, Multi Channel Systems). Spike detection was performed with the MC_rack software (Multi Channel Systems), and the results were further processed with Neuroexplorer (Neuronexus) and MATLAB (MathWorks). Firing rates were measured from 100 ms to 30 ms before the light onset and between 40 ms and 300 ms from the pulse light onset. The first window corresponds to the detection of baseline activity, while the second window corresponds to the detection of the network-mediated response of retinal ganglion cells. In each window the average firing rate was obtained by averaging the instantaneous firing rates from 7 bins around the bin with the maximum firing rate within the window.

### Optical safety

Retinal damage upon light exposure can occur because of three main factors: photo-thermal damage, photo-chemical damage, and thermo-acoustic damage ^19,31^. For 10-ms light pulses delivered at 20 Hz, the MPE could be controlled by the photo-thermal (MPE_T_) or photo-chemical damage (MPE_C_), according to equations (1) and (2), respectively.

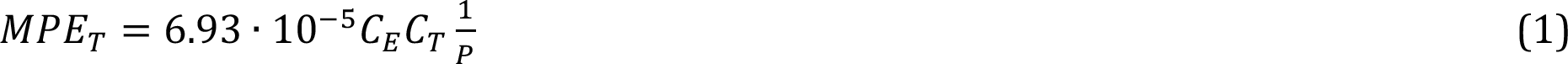

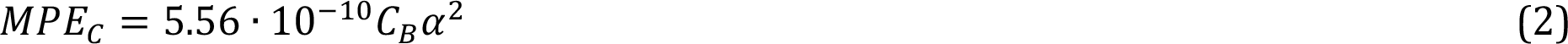

For the POLYRETINA prosthesis, the visual angle of is α = 808.12 mrad with an exposed area of 144.22 mm^2^ ^12^. For λ = 565 nm, both limits apply and *C*_*E*_ = 6.67 . 10^−3^ *α*^2^ *C*_*T*_ = = 1; *P* = 5.44; *C*_*B*_ = 10^0.02(λ−450)^. The limits are MPE_T_ = 55.49 mW and MPE_C_ = 72.45 mW. Therefore, the limiting factor is MPE_T_ which results in 55.49 mW and corresponds to 384.75 μW mm^−2^ for an exposed area of 144.22 mm^2^. For λ = 730 nm and 780 nm, only the MPE_T_ applies and *C*_*E*_ = 6.67 · 10^−3^*α*^2^; *C*_*T*_ = 10^0.002(λ−700)^; *P* = 1; *C*_*B*_ = 1000. Therefore, for λ = 730 nm the MPE_T_ is 346.59 mW, which corresponds to 2.40 mW mm^−2^ for an exposed area of 144.22 mm^2^. For λ = 780 nm, the MPE_T_ is 436.33 mW, which corresponds to 3.03 mW mm^−2^ for an exposed area of 144.22 mm^2^. Because photovoltaic prostheses operate with pulsed light (i.e., in our case 10-ms pulses at 20 Hz of repetition rate), the MPE is increased by a factor of 5 to 1.92, 12.00, and 15.15 mW mm^−2^ respectively for 565, 730, and 780 nm.

### Statistical analysis and graphical representation

Statistical analysis and graphical representation were performed with Prism (GraphPad Software Inc.). The normality test (D’Agostino & Pearson omnibus normality test) was performed in each dataset to justify the use of a parametric or non-parametric test. In each figure p-values were represented as: * p < 0.05, ** p < 0.01, *** p < 0.001, and **** p < 0.0001.

